# The emergent Yo-yo movement of nuclei driven by collective cytoskeletal remodeling in pseudo-synchronous mitotic cycles

**DOI:** 10.1101/662965

**Authors:** Zhiyi Lv, Jan Rosenbaum, Stephan Mohr, Xiaozhu Zhang, Deqing Kong, Helen Preiß, Sebastian Kruss, Karen Alim, Timo Aspelmeier, Jörg Großhans

## Abstract

Many aspects in tissue morphogenesis are attributed to the collective behavior of the participating cells. Yet, the mechanism for emergence of dynamic tissue behavior is not understood completely. Here we report the “yo-yo”-like nuclear drift movement in *Drosophila* syncytial embryo displays typical emergent feature of collective behavior, which is associated with pseudo-synchronous nuclear division cycle. We uncover the direct correlation between the degree of asynchrony of mitosis and the nuclear collective movement. Based on experimental manipulations and numerical simulations, we find the ensemble of spindle elongation, rather than a nucleus’ own spindle, is the main driving force for its drift movement. The cortical F-actin acts as viscoelastic medium to dampen the movements and plays a critical role in restoring the nuclear positions after a mitosis cycle. Our study provides insights into how the interactions between cytoskeleton as individual elements leads to collective movement of the nuclear array on a macroscopic scale.

## Introduction

Collective behaviors emerge from many interacting individuals in the absence of central coordination and supervision. Birds flocks, fish schools, as well as bacterial colonies exhibit spontaneous synchronization and many other complex dynamic functional patterns originating from simple local interaction rules [1–3]. Within a single organism, self-organized collective behaviors are an inseparable part in maintaining its basic biological functions by efficiently driving numerous complex biological processes without external regulations such as neural activation in brain [4–6] and morphogenesis during embryonic development [7,8]. On subcellular scale, actin and microtubules form collective motion in the presence of motor proteins and ATP [9–11]. Yet, despite the ubiquitousness and the significance of emergent collective behavior in biology, our knowledge of the fundamental mechanism of functional collective behaviors are very limited. The major obstacle lies in the typically extremely complex interactions between the individuals in most biological collective behaviors. However, in syncytial insect embryo only direct cytoskeletal interactions exist, owing to the lack of membranes separating the nuclei, which enables dissecting and understanding how the interactions of individuals drive the formation of emergent features at tissue level.

To obtain insights into the molecular mechanism leading to the emergence of a collective flow movement, we study the dynamics of nuclear array in *Drosophila* syncytial blastoderm, where the direct interactions between individuals lead to at least three features of emergent collective behavior observed at tissue level. Firstly, the nuclei divide synchronously to their immediate neighbors but asynchronously to more distant nuclei. This leads to a wave front of mitosis sweeping over the embryo [12,13]. Secondly, the nuclei arrange in an ordered array in interphase following disturbance during nuclear division. Ordering involves interactions by the microtubule asters but also with F-actin at the cortex [14,15]. Thirdly, the nuclei and the cytoplasm undergo stereotypic flow movements following the mitotic wave, which is reminiscent of the toy “yo-yo”. This arises the questions: how does the flow emerge and what are the underlying driving forces? Is there any physiological function of this flow?

The large-scale collective movements are driven by active elements of the cytoskeleton. For example, kinesin-1 and microtubules drive the cytoplasmic streaming during *Drosophila* oogenesis [16]. The cytoplasmic streaming in plant and algae is driven by myosins moving on F-actin [17,18]. Apical constriction mediated by actomyosin can generate the cytoplasmic flow, which compels nuclear spreading in *Drosophila* pre-blastoderm [19] and cell elongation in gastrulation [20].

In *Drosophila* syncytial blastoderm, the nuclei and their associated centrosomes and microtubule asters form an extended two-dimensional array [14]. The dynamics of this array is dominated by two different interactions [21]. Lateral interactions between the neighbor nuclei and centrosomes are mediated by microtubules and the associated motor proteins. The interactions between centrosomes and actin cortex constitute the cortical interactions. How these interactions lead to the stereotypic nuclear movement is unknow.

Beside the force generating mechanism, the material properties of the embryos [22–24] may influence the movement of the nuclei. The actin cytoskeleton inhibits short time-scale movements [21] and promotes ordering of the nuclear array [14]. The actin cortex may act as a viscoelastic medium, to which the centrosomes and their associated nuclei are connected.

Here we found that the nuclear movement was isotropic for individual spindles but anisotropic for the collective flow over several nuclear diameters away from the mitotic wave front and back to the original position slightly later. We comprehensively quantified nuclear trajectories in wild type and mutant embryos and modelled the process by computational simulation. In this way, we uncovered that the ensemble of spindle elongation, rather than a nucleus’ own spindle, is the main driving force for its drift movement. In addition, we defined a function of cortical F-actin for the apparent viscoelastic material properties to restore the nuclear position. Lastly, based on a simulation, we proposed that such nuclear movement is prerequisite for nuclear division in high density in Drosophila syncytial blastoderm.

## Results

### Collective flow and density changes follow the mitotic wave front of pseudo-synchronous nuclear cycles

To elucidate the mechanisms that arise the emergent features of nuclear array dynamics in *Drosophila* syncytial cleavage cycle, we first documented nuclear division cycles using time-lapse microscopy. The nuclei divide slightly asynchronous especially during the last syncytial division in NC13 with a time lag of up to minutes, which is easily visible as a mitotic wave front sweeping over the embryo (Fig. 1A, Supp. Data Fig. S1A, Movie 1). The mitotic wave is driven by Cdk1 activity wave and the molecular processes of the how the chemical wave propagates were dissected [12,25].

**Figure 1.**
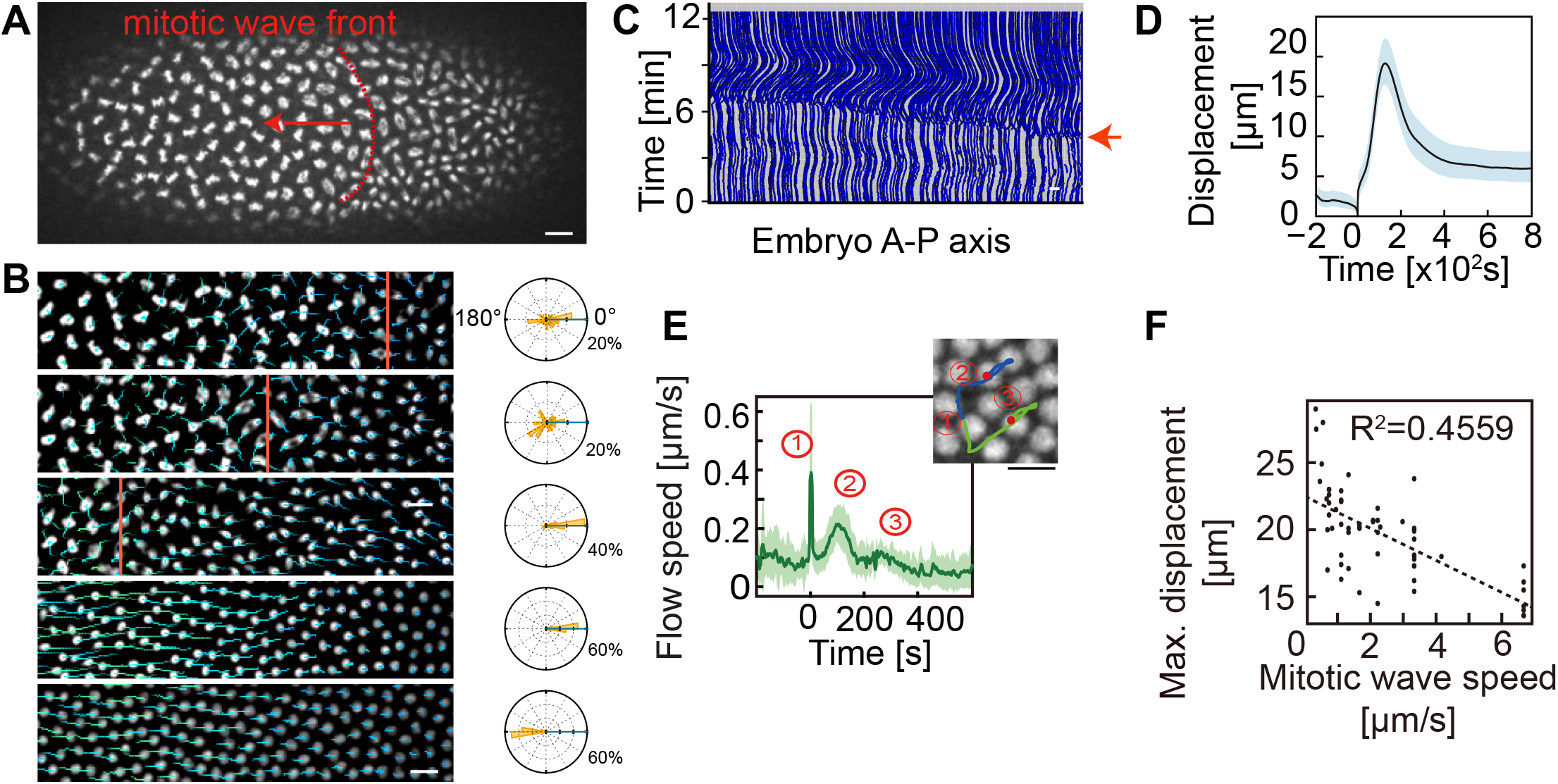
Quantitative assay for nuclear movement. (A), Live image of an embryo during mitosis. The left half is in metaphase, the right half in anaphase. The dotted line in red indicates metaphase-anaphase transition sweeping over the embryo from right to left as a wave front. (B), Snapshots show the nuclear motion. The red line indicates the metaphase-anaphase transition. Blue line is the forward trajectories of nuclei. The right panel is the corresponding angle distribution of nuclear motion. (C), The projection of the nuclear motion. Red arrow indicates the nuclear division. (D, E), Time course of nuclear displacement with the position of the mother nucleus and nuclear speeds during metaphase-anaphase transition as a reference (t=0) (n=260 nuclei in one embryo, representative for all embryos which have been checked). The numbers indicate ①chromosome segregation, ②forth movement away and ③ back movement toward the mitotic wave front. The right panel is a snapshot of nuclear movement labelled with three numbers. The image is the end of the nuclear movement. (F), The maximal displacement plotted against the corresponding speed of the mitotic wave (n=50 embryos).

The mitotic pseudo-synchrony and its corresponding wave front are associated with a stereotypic nuclear movement, which can be readily observed in time lapse movies (Fig. 1B, C, Supp. Data Movie 2). To obtain a precise description of the nuclear movements, we determined the trajectories of all nuclei within the field of view from time lapse recordings of fluorescently labeled nuclei. From the trajectories, we extracted time courses for displacements, velocities, and nuclear density (Fig. 1D, E, Supp. Data Fig. S1B, C). Every nucleus is assigned an individual time axis with the splitting of daughter chromosomes (metaphase-anaphase transition) as reference time t=0.

Concerning displacement (Fig. 1D), the nuclei moved in average about 20 μm which corresponds to about 4–5 nuclear diameters away from the position of their mother nucleus at t=0. The maximal displacement was reached after about 2 min (Fig. 1D). Following maximal displacement, the nuclei then returned to almost their initial position. This movement resembles yo-yo ball, and hereafter we refer it as yo-yo movement. We calculated the speed of nuclear movement from the individual trajectories as the derivative of the trajectories. The averaged flow speed revealed three peaks (Fig. 1E). The first peak corresponds to the chromosome segregation in anaphase with about 0.4 μm/s. The second peak after about 1 to 2 min corresponds to the flow away from the mitotic wave front. The least pronounced, third peak corresponds to the return movement after about 4 min.

Asynchronicity of mitosis might be prerequisite for yo-yo flow. To test whether there is a correlation, we collected data from 50 embryos (Fig. S1A) for quantification of the speed of wave front and maximal displacement. Plotting corresponding parameter sets revealed a negative correlation (R^2^=45%, Fig. 1F), which suggests a slow wave is associated with a large displacement.

As flow is linked to density changes, we next established spatial and temporal maps of nuclear density. Each nucleus was assigned an area and corresponding density according to Voronoi segmentation (Supp. Data Fig. S1B, Movie 3). In the case of synchronous divisions, the density would be expected to double at t=0 (metaphase anaphase transition) and remain constant throughout interphase. In contrast, but consistent with the observed nuclear flow, our measurements revealed a peculiar time of the density. Although initially doubling, the density dropped in telophase before finally reaching the doubled density again a few minutes later (Supp. Data Fig. S1B, C). Corresponding profiles for displacement and flows were detected in preceding nuclear cycles 11 and 12, although in a less pronounced manner (Supp. Data Fig. S2).

The forth and back movement of the nuclei is reminiscent of a spring (Supp. Data Fig. S1D). To obtain a phenomenological description of this behavior, we applied a simple mechanical model to the nuclear trajectories during the period of maximal displacement. By fitting a square function to the displacement curve, we obtained an apparent spring constant for each nucleus (Supp. Data Fig. S1E). The actual value of the apparent spring constant is not informative, since our model and assumptions are too simple. Friction is not included, for example. Yet, the apparent constant helps to compare experimental conditions and mutant phenotypes.

### Isotropic individual behavior is associated with an anisotropic collective flow

Our analysis revealed a collective directional flow of the nuclear array. Yet the individual spindles are isotropically oriented. The axes between daughter nuclei are uniformly distributed over the angles against the anterior-posterior axis of the embryo (Fig. 2A–C). In contrast, the same nuclei almost unidirectionally moved along the embryonic axis a minute later, indicating an anisotropic behavior (Fig. 2A–C). The transition of isotropic individual to anisotropic collective behavior is strikingly obvious in the extreme cases of spindle orientation. In the case of a spindle oriented in parallel to the embryonic axis, one set of chromosomes segregated towards, whereas the daughter chromosomes moved away the wave front during anaphase. One minute later both nuclei moved away from the wave front (Fig. 2D). Similarly, in case of a perpendicular orientation of chromosome segregation, the movement during anaphase was perpendicular to the embryonic axis but along the axis a minute later during collective flow. (Fig. 2D).

**Figure 2.**
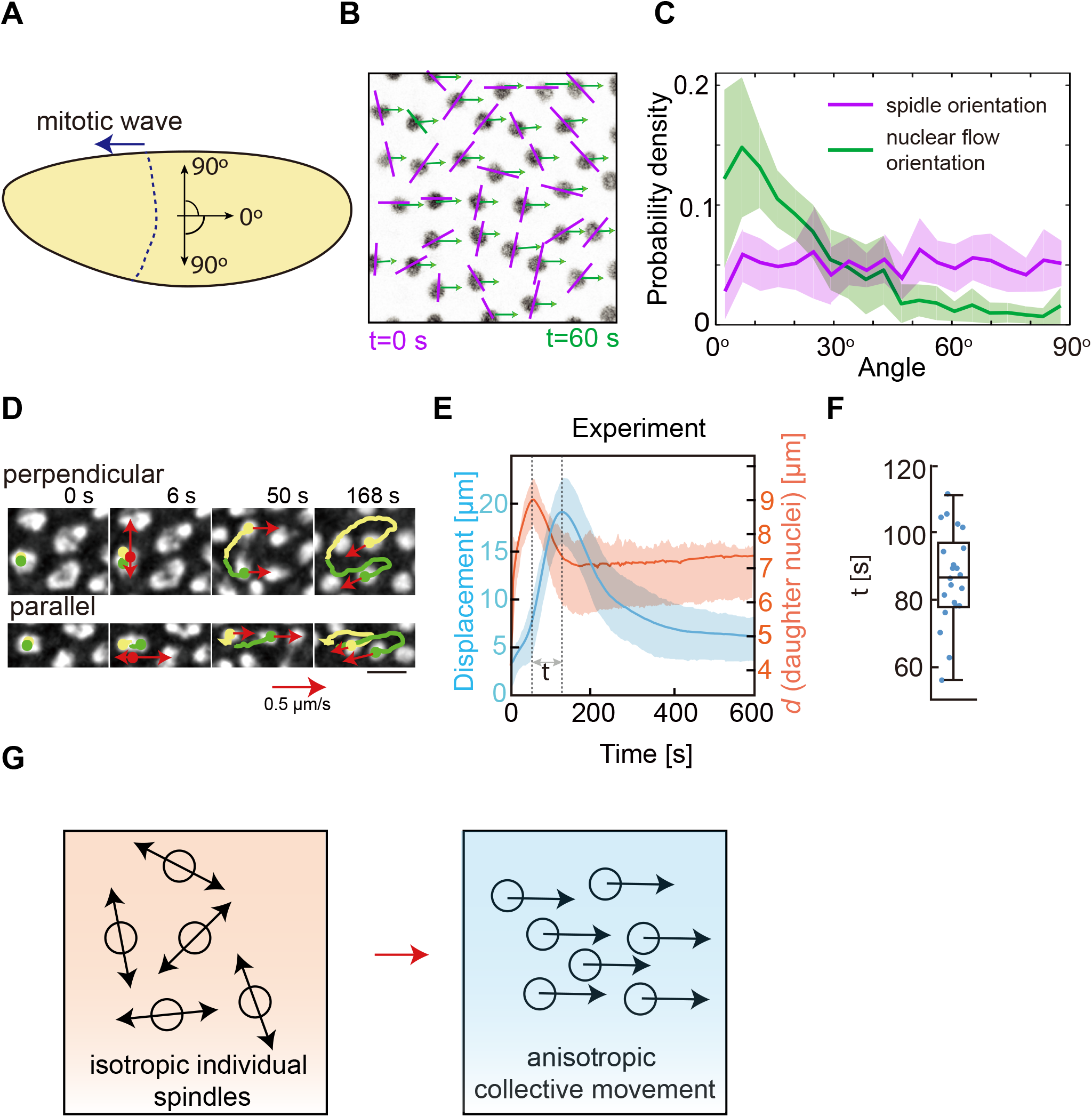
Emergence of collective nuclear movement. (A), Schematic drawing of an embryo with definition of angles. (B), Image from live imaging after mitosis. Orientations of the corresponding spindle at t=0 s and the directions of nuclear movement at t = 60 s are indicated by a magenta bar and green arrow, respectively. (C), Distribution of observed angles for spindle orientation at t=0 s and nuclear movement at t=60 s. (n=20 embryos including 6230 nuclei). (D), Image series with previous trajectories showing cases of perpendicular and parallel mitosis. (E), Time course of nuclear displacement (blue) and distance between corresponding daughter nuclei (orange). The time lag (τ) between maxima is indicated (n=260 nuclei in one embryo, representative for all embryos we have checked). (F), Distribution of the time lag (τ) (n=21 embryos). (G), The schematic drawing of the transition from individual movement to collective behavior. Data are mean±s.e.m. Scale bar: 10 μm

The uncoupling of spindle behavior and nuclear behavior was seen not only in the orientation of the movements, but also in the time course of the two behaviors. The maximal displacement was reached only about 1.5 min after the maximal distance between daughter nuclei (maximal spindle length) was reached (Fig. 2E, F), suggesting the maximal displacement was achieved when the nuclei are in telophase. These findings demonstrated that these two processes were mechanistically not directly linked at the individual level, since spindles were isotropically oriented and preceded the flow behavior (Fig. 2G).

During collective flow the nuclei may move as individuals characterized by neighbor exchanges. Alternatively, nuclei may behave as an array, which would be indicated by fixed neighbor relationships. To distinguish these options, we labeled groups of cells before mitosis and followed them during the course of chromosome segregation and collective flow. We found that the nuclei moved as an array. The groups of nuclei did neither intersperse with unlabeled nuclei nor nuclei of the other groups indicating that neighbor relationships were maintained during mitosis and collective flow despite the movement over several nuclear diameters (Supp. Data Fig. S3A). In addition, we measured the displacement fields of nuclei using a particle image velocity algorithm. We found that the nuclei motion is similar to a laminar flow (Supp. Data Fig. S3B, Movie 4). In summary, our observations indicate that the nuclear layer phenomenologically behaves like an elastic sheet with fixed neighbor relationships.

### Computational modeling of nuclear movement

To gain a better understanding for how the isotropic spindles gives rise to an anisotropic collective flow, we conducted computational simulations. Starting from a computer model for static nuclear interactions in interphase [15], we added a time axis for the interactions. The model is based on active and passive forces (Fig. 3A). Stochastic active forces repulse adjacent nuclei, thus resembling the sliding activity of motor proteins, e. g. Kinesin-5, on antiparallel aligned microtubules. In addition, a passive elastic force leads to repulsion accounting for the embedding of the nuclei into the cytoplasm and cytoskeleton. This may include the link of the nuclei to the cortex. Chromosome segregation is triggered at t=0 by a separation force acting between the daughter nuclei. The interaction forces are dynamic according to the mitotic stage and interphase (Supp. Data Fig. S4). For example, the active force is low in anaphase, since astral microtubules prominently appear only in telo- and interphase. Similarly, the passive force increases in telo- and interphase as cortical actin increases during these stages. The segregation force decays in telo and interphase. Balancing the magnitude of passive and active forces over time, model simulation reproduced the experimentally observed stereotypic flow behavior (Fig. 3B, C, Supp. Data Movie 5).

**Figure 3.**
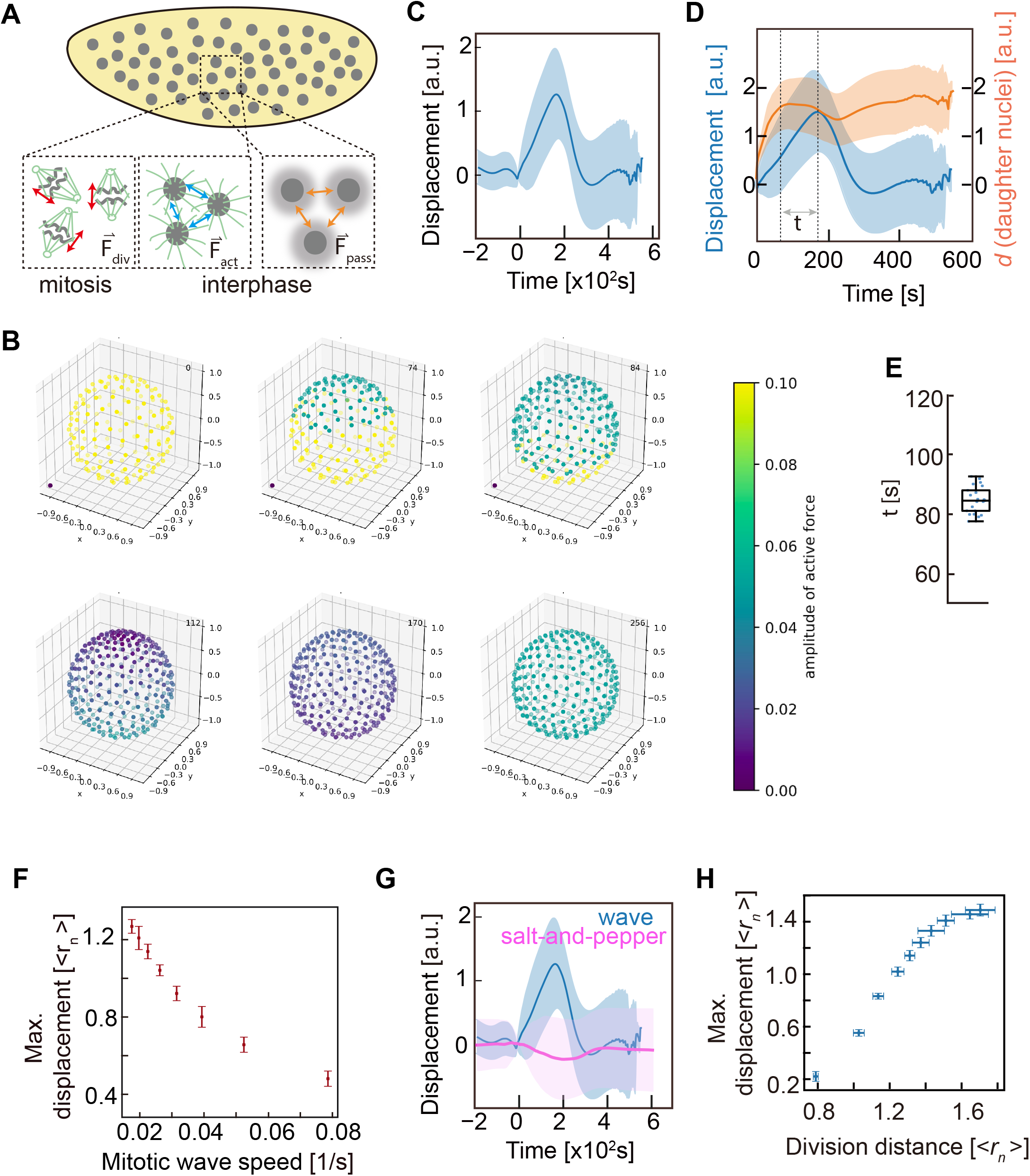
Numerical simulation of nuclear movement. (A), Scheme of division, active and passive forces in syncytial embryo. (B), Snapshots from the simulation. Nuclei were projections. Color code indicates speed of nuclear movement. (C), Time course of nuclear displacement (n=300 nuclei). (D), Time course of nuclear displacement (blue) and distance between corresponding daughter nuclei (orange) (n=300 nuclei). The time lag (τ) between maxima is indicated. (E), Distribution of the time lag (τ) (n=21 embryos). (F), The negative correlation between maximal displacement and the corresponding speed of the mitotic wave. (G), Time course of nuclear displacement in wave (wild type) embryo and “salt-and-pepper pattern” embryo (n=300 nuclei in both cases, respectively). (H), The positive correlation between maximal displacement and division distance. Data are mean±s.e.m.

Strikingly, the simulations reproduce other features of nuclear yo-yo movement. Firstly, maximal spindle length precedes maximal displacement with a time lag of 1.5 min (Fig. 3D, E vs Fig. 2E, F). Secondly, the speed of the mitotic wave front is negatively correlated with the nuclear maximal displacement (Fig. 3F vs Fig. 1F). In addition, the simulations predict no collective movement of nuclei with asynchronous nuclear divisions in a “salt-and-pepper” pattern (Fig. 3G). Importantly, the simulations predict that the force for separation of the daughter nuclei positively correlates with maximal distance between daughter nuclei and importantly with maximal displacement (Fig. 3H). Thus, our simulations predict that spindle elongation is a major driving force for nuclear movement. Mechanistically, this is not a simple relationship because spindles are isotropically oriented, whereas the direction of displacement is anisotropic.

### Ensemble spindle elongation is the driving force for nuclear displacement

To test the prediction that spindle elongation drives the yo-yo movement, we preformed the laser cutting on single spindle. However, the spindle recovered in second-scale and no effects on nuclear motion (Supp. Data Fig. S5). To circumvent this problem, we developed a method to reduce the spindle length globally, without hampering the nuclear separation. Spindle elongation in anaphase B requires the four-headed microtubule motor Kinesin 5, which can slide microtubules against each other [26]. We employed embryos, in which endogenous Kinesin 5 was substituted by a version susceptible to TEV protease [27]. We titrated the amount of TEV protease to achieve a partial depletion which still allowed completion of mitosis. Spindles in these embryos were short (Fig. 4A). The average maximal spindle length was 8 μm instead of 10 μm in wild type embryos. Complementary, we employed embryos from females homozygous for *Map60*, which also displayed short spindles [14] with an average length of 9 μm (Fig. 4A). Quantification of nuclear movement revealed a strongly reduced maximal displacement in both experimental conditions and thus a positive correlation of spindle length and maximal displacement (Fig. 4B, C, D). In summary, analysis of mutant embryos with shorter division distance support an ensemble spindle elongation constitutes a major driving force for the nuclear movement thereafter.

**Figure 4.**
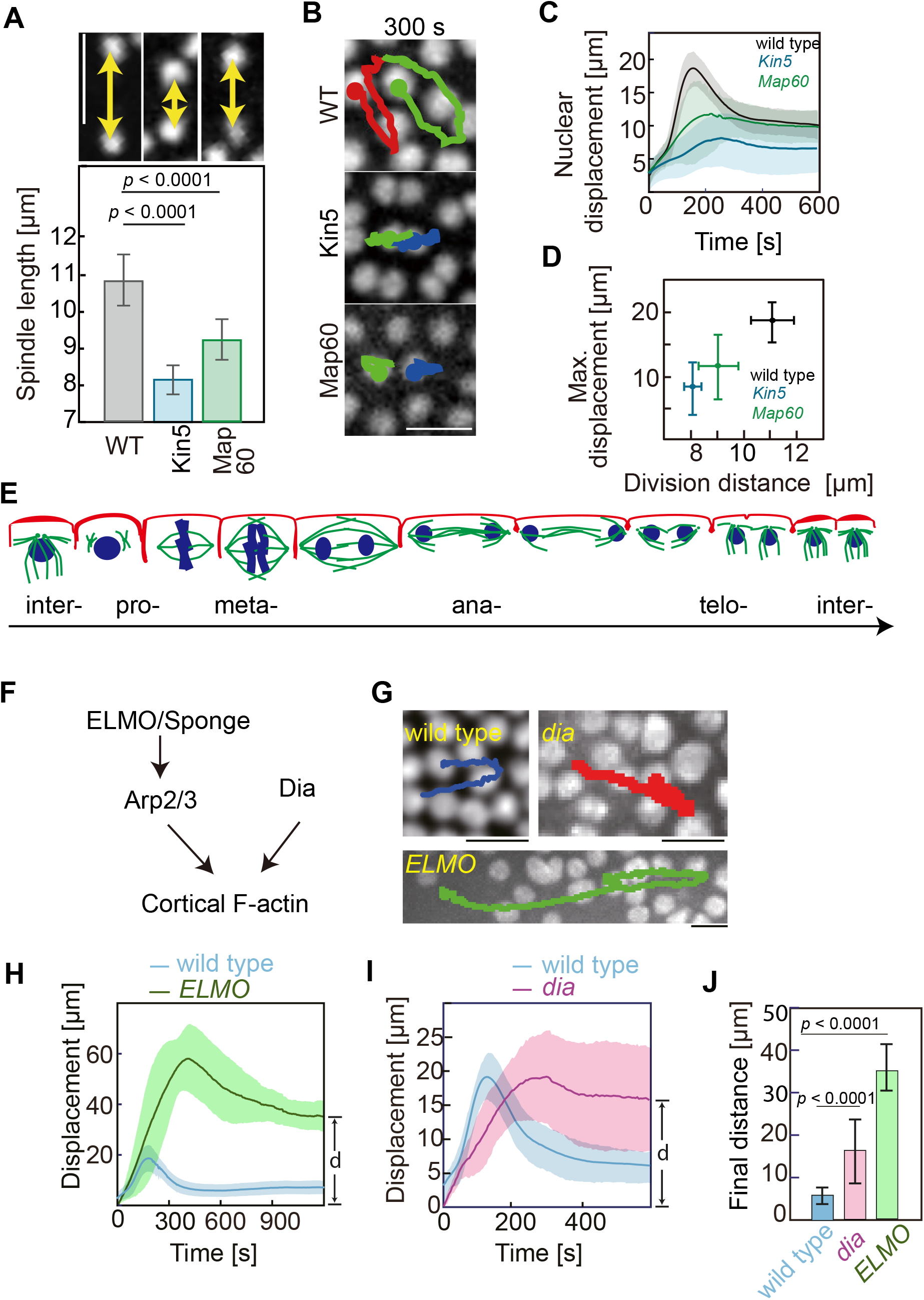
Forces mediating Yo-Yo movement of nuclei *in vivo*. (A). Spindle length/distance between respective daughter nuclei in embryos partially depleted of Kinesin-5 and *Map60* mutants (n=30 spindle in 3 embryos for each genotype). (B), Images from live image. The trajectories of two nuclei over 300 s were plotted into the images. Images were the end of the movement. (C), Time course of nuclear displacement. (D), Maximal displacement was plotted against corresponding division distance. (n=290 nuclei in wild type, 256 nuclei in *kin5*, 292 nuclei in *Map60* in Fig. C and D). (E), Scheme of cortical actin dynamics during nuclear division cycles. (F), ELMO and Dia are involved in F-actin cortex formation. (G), Images from movies of *dia* and *ELMO* mutants. Nuclear trajectories over 10 min are plotted into the images. Images were the end of the nuclear movement. (H, J), Time course of displacement in *ELMO* and *dia* mutants. (J), The final distance between the current position and original position of the nuclei after yo-yo movement, as indicated as “d” in H and I (n=260 nuclei in wild type, 45 nuclei in *ELMO* mutant and 90 in *dia* mutant embryo in Fig. H, I, J). The Data are mean±s.e.m. Scale bar: 10 μm.

### F-actin cortex is required for the return movement

The simulation predicts a function of passive force in yo-yo movement. Cortical F-actin is a promising candidate. It has been previously reported that the nuclei are strongly attached to F-actin cortex [28]. The actin cortex suppresses the fluctuation movements of centrosomes [21] and contributes to an ordered nuclear array in interphase [14]. Cortical F-actin undergoes stereotypic remodeling during the course of nuclear cycles [29] (Fig. 4E, Supp. Data Fig. S6A). We quantified total F-actin with a Utrophin-GFP as a marker [30]. We found that the signal dropped in mitosis, with lowest level during anaphase and steadily increased afterwards (Supp. Data Fig. S6A, B). Given the timing of this dynamics, it is conceivable that cortical actin plays an important part in controlling nuclear movement.

To test this conceivable function of the cortical F-actin, we employed two mutants to genetically interfere with the organization of the actin cortex (Fig4. F, Supp. Data Fig. S6G). Firstly, we prevented the formation of actin caps with the mutant *ELMO* [21,31]. ELMO forms part of an unconventional guanyl nucleotide exchange factor, which activates Rac signaling in a complex with Sponge/DOCK [32]. *ELMO* mutants lack any actin caps and are characterized by a uniformly structured cortical F-actin [21] (Supp. Data Fig. S6G). Secondly, we employed *dia* mutants [33–35]. Dia is a founding member of the formin family, which nucleate and polymerize linear actin filaments. *dia* mutants lack metaphase furrows but contain actin caps [33] (Supp. Data Fig. S6G).

We applied our quantitative assay to *ELMO* embryos. A strongly increased nuclear mobility was obvious in time lapse movies (Fig. 4G). Quantification of nuclear trajectories revealed a maximal nuclear displacement of 60 μm as compared to 20 μm in wild type embryos (Fig. 4H, Supp. Data Fig. S6H). In addition to the threefold increased displacement, we observed as a second phenotype that the nuclei did not return to their initial position in *ELMO* mutants (Fig. 4G, H). The impaired back movement indicates a loss of the spring-like behavior. Consistently, we calculated an almost 10-fold reduced spring constant at the turning point of the nuclear trajectories (Supp. Data Fig. S6I). A similar behavior and profiles were detected in NC12 of ELMO embryos (Supp. Data Fig. S6J, K).

We also detected changes in nuclear movement in *dia* mutants. Similar to *ELMO* mutants, we observed a loss of the spring-like back movement. The nuclei did not return to their initial position and the spring constant was almost 10-fold reduced (Fig. 4I, J, Supp. Data Fig. S6H, I). In contrast to *ELMO*, the maximal displacement was similar to wild type indicating that the stabilizing/viscos function of the cortex does not depend on *dia*. The neighborhood relationships were largely maintained in *dia* and *ELMO* embryos (Supp. Data Fig. S7). In summary, by employing two mutants affecting F-actin organization, we identified distinct functions of the actin cortex. F-actin is required for the back movement as revealed by the reduced apparent spring constant and the permanent displacement. The *ELMO*-dependent organization into caps appears to be important for limiting nuclear movement.

### Long mitotic spindles and distance between daughter nuclei require pseudo-synchronous nuclear cycles

To obtain the further insight into the mitotic wave and consequent yo-yo movement in syncytial embryo, we documented the nuclear separation profile. We found that in mitosis the daughter nuclei are separated by an overshooting spindle, which pushes apart the daughter nuclei more than the average inter-nuclear distance (Fig. 5A, B). With every division during the syncytial blastoderm (nuclear cycles NC 10–13), the nuclear density at the cortex doubles (Supp. Data Fig. S8). The nuclei divide in a wave manner especially during the last syncytial division. The mitotic wave is driven by Cdk1 activity wave, and the activity of DNA replication checkpoint is important in the slowdown of the wave, which occurs in later cycles [12,25]. However, the function of this pseudo-synchrony and the consequent yo-yo movement is unknown. Given the increase of asynchrony with nuclear density, we speculated that spindle overshooting with nuclear crowding may pose a problem for synchronous divisions.

**Figure 5.**
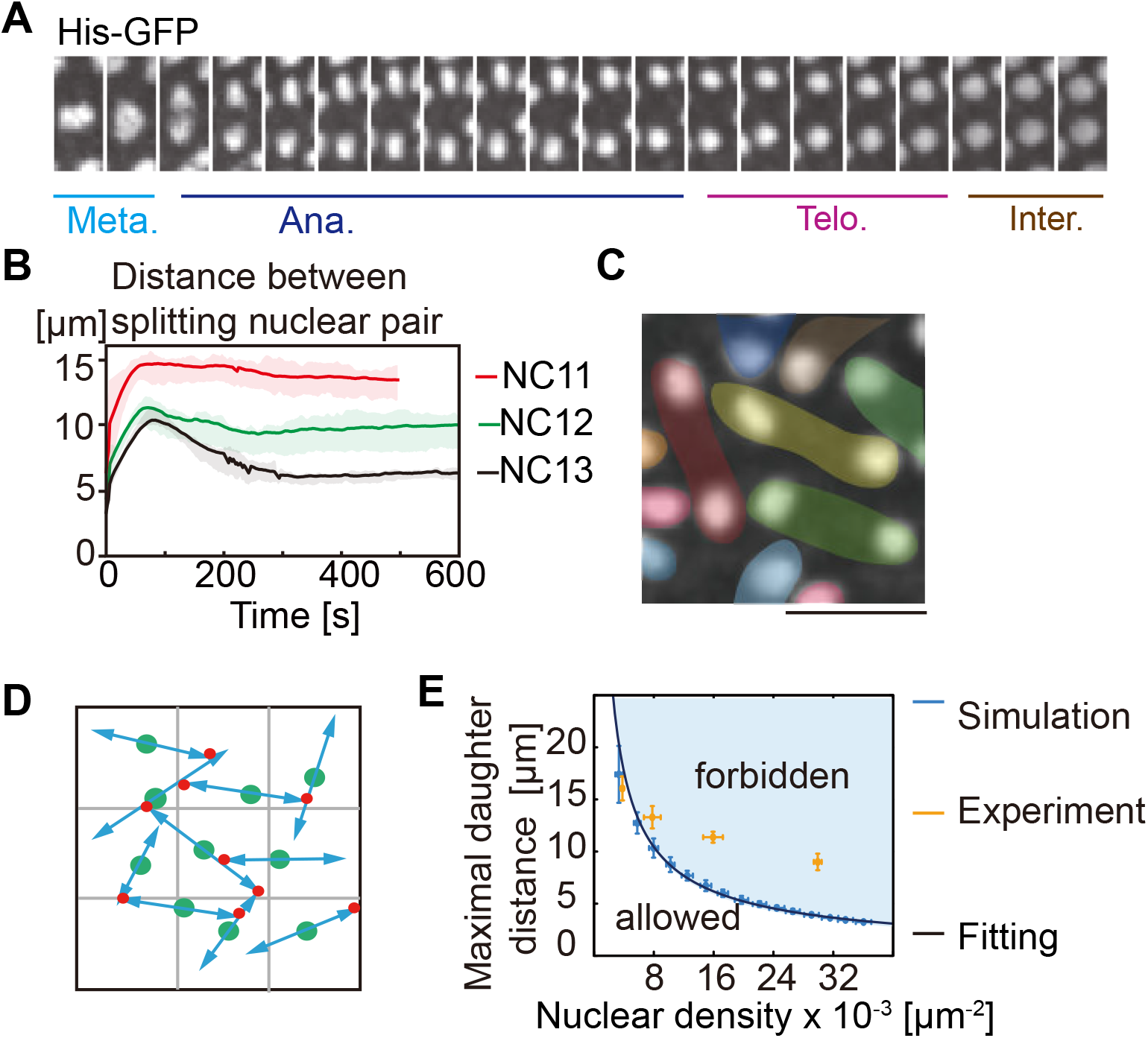
Long mitotic spindles require pseudo-synchronous nuclear cycles. (A), Live image series of the process of nuclear dividing. (B), Quantification of the distance between splitting nuclear pair in cleavage cycles. (C) The protected area between daughter nuclei containing the spindle is indicated by an overlaid colored area. (D), Illustration of the simulation. Green dots indicate nuclei. Blue arrows indicate spindles. Red dots indicate the touching of the neighbor spindles, leading to the spindle growth stops. (E), Spindle lengths/distances between daughter nuclei are plotted against the corresponding nuclear density. Data from simulation (blue) and measurements in embryos in NC11, NC12, NC13 in diploids and NC14 in haploids (orange) (n=15 spindles in each embryo and 3 embryos for each type).

To test this hypothesis, we employed the *mei41 zld* double mutant embryos [36], as well as the *grp nmk* double mutant embryos [12], to experimentally reduce the nuclear division time in cycle 13. We planned to check how much the spindles were able to elongate in a similar level of nuclear crowding as in wild type embryo, but with a more synchronous division manner. Unfortunately, the chromosomes were not separated at cycle 13 in these mutant embryos (Supp. Data Fig. S9). To circumvent this problem, we simulated nuclear divisions within a limited area. We assigned each nucleus and mitotic spindles a protected area representing the entity in real embryos (Fig. 5C). After chromosome division, the daughter nuclei were pushed apart until reaching the protected area of a neighboring spindles/pair of daughter nuclei, thus assessing the maximal possible distance between daughter nuclei (maximal spindle length). We assumed synchronous divisions and symmetric spindles with isotropic orientations (Fig. 5D). Our simulations showed that the maximal spindle length decreased with an increase in nuclear number and thus marked the transition line between structurally allowed and forbidden regime of combinations of nuclear density and spindle length (Fig. 5E). Next, we measured the maximal distance between daughter nuclei and their corresponding nuclear densities in wild type embryos. We also included data from haploid embryos, which undergo an extra nuclear division. Only the parameters of NC11 fell into the allowed area, whereas the parameters of NC12, NC13, and NC14 in haploids fell into the forbidden regime. This analysis indicated that a synchronous division with the observed spindle lengths was possible only in the early cycles but impossible in later cycles. Thus, pseudo-synchrony allows for the observed spindle length in NC12, NC13 and NC14 in haploid embryos.

## Discussion

The direct interactions between the nuclei and their associated cytoskeleton are a special feature of syncytial embryos. Due to the lack of separating cell membranes, microtubule asters originating from the centrosomes associated with each nucleus form an extended network of hundreds to thousands of elements. Emergent features arise in this network by summing up the behavior of individual elements, such as fluctuations or duplication, and their interactions, such as repulsion. The analysis of the mechanism underlying the emergent features is essential for understanding how the individual cells function collectively to form a tissue.

We identified an anisotropic flow of the nuclear array as an emergent feature. Based on a morphodynamical analysis of the nuclear array in wild type and mutant embryos together with computational simulations, we analyzed the mechanism of the flow behavior. In this way we identified spindle elongation drives nuclei moving away whereas cortical F-actin restores the nuclear positions, which is necessary for the following development. The emergent nature of the nuclear flow becomes obvious, since individual behavior is strikingly different than the collective behavior of the nuclear array. Nuclei divide with an isotropic orientation, whereas the flow direction is anisotropic. Furthermore, the maximal division distance is about 10 μm, whereas the maximal displacement is about 20 μm.

In addition to the driving force of the nuclear yo-yo movement, our “limited area” simulation (Fig. 5C, D, E) demonstrates the necessity of such movement. To complete the nuclear division in a limited space with high nuclear density, two strategies could be utilized. The first strategy is to reduce the spindle length, as what happens in *Map60* mutant and Kinesin-5 partially depleted embryos. This might raise the risk that the genetic materials cannot separate completely. The second strategy is that nuclei divide in an asynchronous manner. Besides *Drosophila*, the nuclear division asynchrony was observed in beetle *Tribolium castaneum* [37], implying this might be a conserved mechanism in insect syncytial embryos. The nuclear directional movement is the consequence of the asynchronous divisions. With strongly synchronous divisions, the pushing forces of mitotic spindles would generate a spatially isotropic force distribution. Consequently, the nuclei would not move due to a balance in forces. However, in the case of pseudo-synchronous divisions, the force balance is broken leading to an imbalance and thus a flow away from the wave front. The repulsive force between daughter nuclei increases in anaphase pushing the daughter nuclei apart, followed by a drop in telophase due to spindle disassembly (Supp. Data Fig. S11A). The summing up of all nuclei in an embryo at a given mitotic time results in an asymmetric force field, which likely determines the directionality of the nuclear flow in telophase (Supp. Data Fig. S11B).

Upon the nuclei reach the maximal displacements, they return to the starting positions. Consistency of nuclear position among different cleavage cycles is important for maintaining the positioning information provided by morphogens. We identified a contribution of the actin cortex to the viscoelastic feature of the nuclear movement, i.e. that nuclei return to their starting position. This is consistent with previous findings that the elasticity of *Drosophila* embryonic cortex in cellularization stage depends on the actin cytoskeleton [23,24]. In addition, the structure of actin cortex is undergoing remodeling, which may contribute to the returning movements by actively changing the cortical material properties in time and space.

Collective behaviors driven by the integration of forces originated from cytoskeletal networks are indispensable in biological systems. Some basic ingredients of the system — the mechanical properties of the cytoskeleton and the function of motor proteins have been well studied *in vitro*. The morphology of early *Drosophila* embryos has also been extensively imaged. However, the assembly of the puzzle to achieve a quantitative understanding of the molecular mechanics behind the dynamical self-organization of the rapidly developing embryo, has only begun to be explored. Our study of the dynamical properties of the syncytial embryo is a first step towards our long-term goal to understand how cells mechanically interact with each other and collectively function as active matter forming a tissue.

## Methods and Materials

### *Drosophila* Genetics

Fly stocks were obtained from the Bloomington Drosophila Stock Center [38,39], unless otherwise noted. Fly strains used in this study are the followings: w; Histone2Av-GFP. *w;* mCherry-Tubulin, Histone2Av-GFP. *w*; sqh-Utr-GFP/CyO; ubi-His2Av-RFP[30]. *Map60*^*KG00506*^. *w*; ubi-GFP-D-Cad *dia*^*5*^ Frt^2L^ ubi-His2Av-RFP/CyO [33]. *w*; *ELMO*^*367*^ Frt^2L^/CyO [21]. *w Hira*^*ssm*^/FM7c, w^a^ B [40]. His2Av-RFP; Kinesin5-[TEV]-GFP [27]. *mei41 zld/*FM7 [36]. *grp nmk*. Sqh-GFP, Histone2Av-GFP. Fly stocks were kept at 25°C on a standard cornmeal food. Germline clones of *dia* and *ELMO* were induced by crossing with corresponding Frt chromosomes and the following heat shock at 37°C for one hour on two consecutive days after hatching.

### Phalloidin staining and imaging

Wild type embryos and embryos from *dia* and *ELMO* germline clones were fixed with 8% formaldehyde according to standard procedures. The vitelline membrane was manually removed. Fixed embryos were incubated with phalloidin-Alexa 488 (1:500, Thermo Fisher) for 1.5 h. After rinsing three times and washing three times for 15 min each with PBT (PBS(Phosphate-Buffered Saline) with 0.1% Tween 20), embryos were stained with DAPI (4′,6-Diamidine-2′-phenylindole dihydrochloride) (0.2 μg/ml) for 10 min, rinsed three times in PBT, washed in PBT for 10 min and mounted in Aqua-Poly/Mount (Polysciences). The images of fixed embryos were acquired using a Zeiss LSM780 confocal microscope.

### Microinjection

1–2 h old embryos were collected, dechorionated with 50% bleach solution for 90 s, rinsed thoroughly with deionized water. After aligning on a coverslip, the embryos were desiccated for 10 min, and covered with halocarbon oil (Voltalef 10S, Lehmann & Voss). TEV protease (a gift from Dirk Görlich) and Histone1-Alexa-488 protein (2 mg/ml, Thermo Fisher) were injected to the desired embryos using Microinjector FemtoJet® (Eppendorf) on an inverted microscope. Short spindle was induced by TEV injection in to the embryos expressing Histone2Av-RFP and Kinesin5-[TEV]-GFP. To get the right concentration of TEV protease for injection, we injected TEV with a serial dilution that covers a range of concentration from 10 μM to 0.1μM. We found 1μM was robust to achieve a partial depletion allowing mitosis but with shorter spindles during the cleavage cycle.

### Live imaging for nuclear dynamics

Nuclear dynamics was recorded by movies of embryos with the fluorescently labeled nuclei, by expression of Histone2Av-GFP or injection of Histone1-Alexa-488 protein. Embryos were attached on a coverslip coated with embryo glue and covered with halocarbon oil. Time-lapse images were recorded on a spinning disc microscope (Zeiss, 25x/NA0.7 multi immersion) with an emCCD camera (Photometrics, Evolve 512). To ensure reliable tracking of the nuclei, the frame rate was 0.5–0.2 Hz with 4 axial sections, covering 8 μm. Images were merged maximal intensity projections (Fiji/ImageJ[41]).

### Images process and quantification

Imaging segmentation and analysis were performed with custom-written Python algorithms. The software code is available on request. Briefly, the nuclear positions were detected as blob-like features of size σ_*i*_ at position (*x*_*i*_, *y*_*i*_) by finding the maxima (*x*_*i*_, *y*_*i*_ σ_*i*_) of a rescaled Laplacian of Gaussian (LoG) function

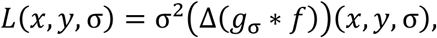

where *f*_*t*_ (*x*, *y*) is the nuclei gray-scale value at time *t*, *g*_σ_ (*x*, *y*) is Gaussian kernel of width σ, and “*g* * *f*” stands for the convolution of function *g* and *f*. When multiple blobs were detected in a single nucleus, we deleted a neighboring blob *b*_2_ of *b*_1_ with a heuristic test function *T*

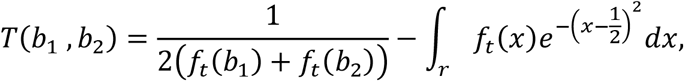

where γ : [0, 1] → ℝ^2^ is the straight line from *b*_1_ of *b*_2_.

We tracked the nuclei across frames based on a proximity criterion. The distance between nucleus *k* in frame *i* and *l* in frame *i* + 1 was defined as

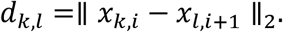

We determined the interval of mitosis time using the *k*-means-clustering algorithm on the observed nucleus positions at time *t*. If a new blob was detected, we considered this nucleus and its nearest neighbor were daughter nuclei from a recent mitosis, and set their internal nucleus clock to 0. Calculations of nuclear displacement, speed, nuclear density, spindle length and orientation were done for each nucleus in its own eigentime after mitosis.

### Laser ablation

Stage embryos expressing Histone2Av-GFP and Cherry-Tubulin were used. Cross-section images were recorded in the Cherry channel with a frame rate of 1/s on a spinning disc microscope (100x/oil, NA1.4) with a CCD camera. Spindle apparatus was ablated at spindle midzone by a line of 355 nm YAG laser (DPSL-355/14, Rapp Opto Electonic) with the 15% of laser power, and around 400 ms exposure time during the recording mode (100x oil, NA 1.4) (Fig. S5).

### Particle Imaging Velocimetry (PIV) analyses

Particle Imaging Velocimetry (PIV) of Histone2Av-GFP images (in Fig. S3, S7) analysis was performed using square interrogation windows of side 16 pixels with an overlap of 10 s “PIVlab” in MATLAB.

### Quantification of F-actin over cell cycle

Embryos expressing Histone2Av-RFP; Utrophin-GFP were imaged with a Zeiss LSM780 confocal microscope (25x/NA0.7 multi immersion). The frame rate was 0.1 Hz, and 10 μm was covered in z direction. Utrophin-GFP stacks were merged by average intensity projection (Fiji/ImageJ). F-actin was quantified manually with Fiji/ImageJ.

### “Limited area” simulation of synchronous mitosis

In the simulation, the nuclei are randomly placed in a 50um*50um square via Poisson-disc sampling, which produces random tightly-packed locations with pair-wise distances not smaller than a specified value d_disc_>4um. We assume the nuclei divide simultaneously and form mitotic spindles with isotropic orientations. As the spindles extend with a constant speed, we check at each time step if any two of the spindles touch each other by scanning a restricted area of 4um*4um in the vicinity of each spindle (see supplementary video). If a spindle touches at least one other, we assume it stops extending and reaches its maximal length due to limited space. When all spindles reach their respective maximal lengths, the simulation is ended and we compute the average maximal length l_max_ over all spindles in the simulation. For each fixed d_disc_, we run the simulation 50 times, producing 50 l_max_ values for various nuclear density around 1/(4d_disc_^2^). The mean of the 50 l_max_ values and the mean of the 50 nuclear densities provide the coordinates of one point in Fig.5E. The x- and y- error bars indicate the respective standard deviations. Varying the minimum distance d_disc_ between nuclei, we obtain the mean l_max_ values for a range of nuclear densities. The data from simulation are fitted to a power-law function (solid curve in Fig. 5E) with the method of least squares.

### Computational modeling of the nuclear movement

We extend the model by Kaiser, et al [15], which has previously been successful in modeling static nuclei ordering during interphase, to now account for nuclei dynamics during mitosis also now incorporating the spherical topology of the embryo: Nuclei, positioned at 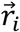, move due to active forces, 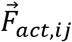, exerted by motor-activated pushing apart of overlapping microtubule asters, and due to passive repulsive, 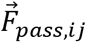, arising from the visco-elastic matrix embedding the nuclei, built mainly from cytoskeletal actin. The overdamped equation of motion is given by 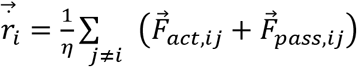, where *η* = 6*πaμ* ≈ 56 × 10^−6^N m^−1^s^−1^ denotes the effective drag coefficient for the approximately circular nuclei [42], where a ≈ 3 μm is the nuclear radius and *μ* ≈1 Pa is the viscosity of the matrix [22,23]. Both forces decay in space following 1 / *r*^4^. For the active force, this is justified because the maximal force a single microtubule can exert scales like 1 / *r*^2^ and the density of microtubuli decays with 1 / *r*^2^ in two dimensions. For a detailed justification on the passive force, see Kaiser, et al [15].

The time dependence of the forces results from time dependent individual force amplitudes contributed by each nucleus. In detail, forces are given by 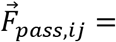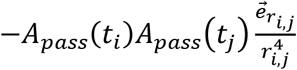 and 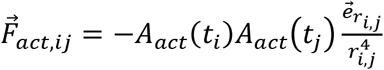, where 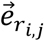 and *r*_*i,j*_ denote the unit vector and the distance between nuclei *i*, *j*, respectively. Nuclei divide when their age reaches *t*_*i*_ = *t*_div_, and ages are initialized as 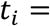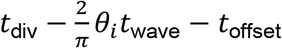, where *θ*_*i*_ is the polar coordinate of nucleus i. This initialization leads to the division wave. For numerical stability, upon division, the two daughter nuclei are placed a short distance *r*_div_ apart with random orientation, their center of mass coinciding with the position of their mother.

To capture the dynamics of cytoskeletal elements during mitosis we subdivided the time course of events into the series: *t*_spindle ass_ < *t*_spindle const_ < *t*_div_ < *t*_sp diss_ < *t*_MT ass_ < *t*_actin ass_ < *t*_MT inter_ < *t*_actin inter_. Passive forces change only between *t*_spindle ass_ < *t*_spindle const_ when actin caps shrink while spindles assemble generating balancing forces and during regrowth of actin caps between *t*_actin ass_ < *t*_actin inter_ before entering interphase. Active forces in contrast are much more dynamic: Between *t*_spindle ass_ < *t*_spindle const_ spindles assemble and the active force grows to exert a maximal force of *h*_spindle_. After division *t*_div_ < *t*_spindle diss_ spindles move daughter nuclei apart: The active force between the division partners is now increased by an additional time-dependent factor *A*_div_ that reaches values of up to 3, 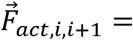 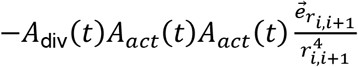, where t is the common time of both daughter nuclei. After *t*_spindle diss_, forces between paired daughter nuclei reduce linearly to regular levels. Meanwhile, for all other nuclei, after division the active force is halved since each nucleus is only associated with one centrosome, instead of two as before. This is balanced by the effective increase of passive forces as nuclei are closer packed due to division while the passive force amplitude stays constant. Between *t*_spindle diss_ < *t*_MT ass_ spindles disassemble before the microtubule asters regrow during *t*_MT ass_ < *t*_MT inter_. All dynamics are interpolated linearly. In detail:

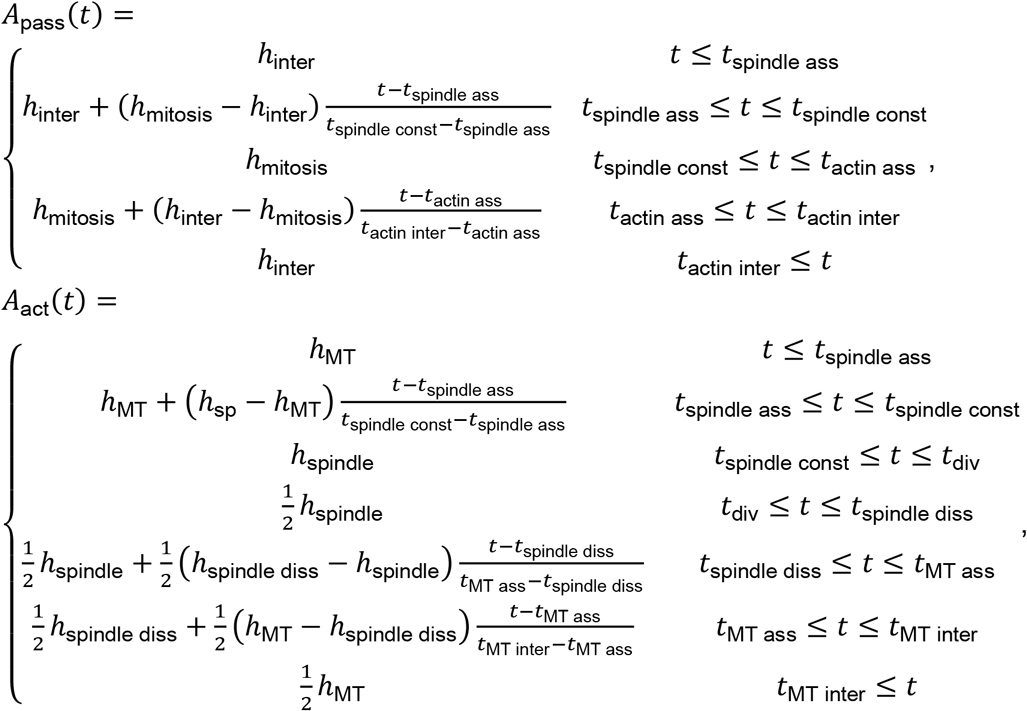

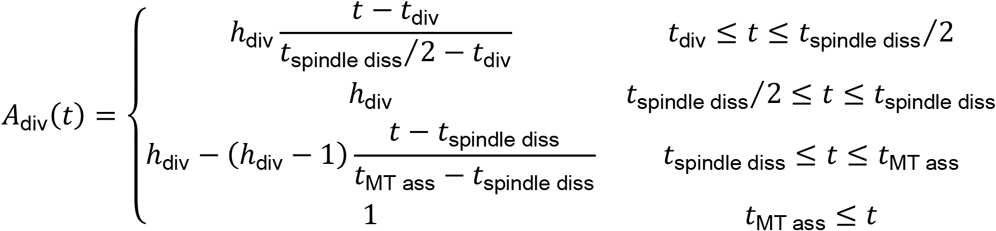

Note that changes in both passive and active forces only get out of balance - initiating nuclei motion - during spindle disassembly with significant time delay relative to the time point of nuclei division. At that point in time nuclei that have not divided yet exert a stronger repulsive force than the already divided nuclei, since their actin caps and microtubule asters are not fully reformed. Therefore, nuclei move toward the region of higher nuclei density, only returning back when actin caps and microtubule asters are forming again.

Choices for the model parameters are found in the table below. Their magnitudes are chosen to match the length of the cell cycle, 800s, and the maximal force exerted by a single microtubule, which is around 3pN according to [43]. In total, the forces on a single nucleus range between 10-100pN, the same order of magnitude as the force applied to a single magnetic microparticle by [24] to move it through the cellularizing tissue in an early drosophila embryo. Note that computing the force involves multiplying the two amplitudes and dividing by the distance to the power of four.

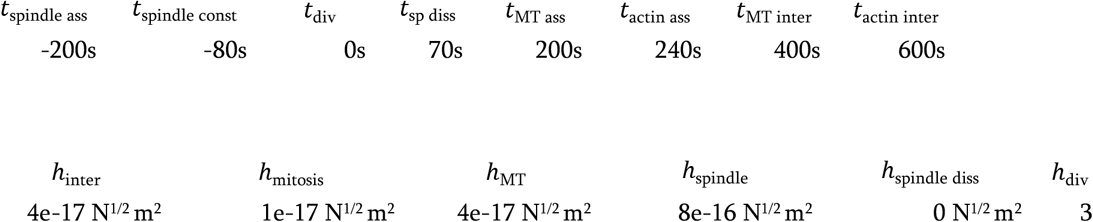

### Spring constant fitting

The data sets consisted of 1–5 nuclear displacement curves for 2–4 embryos of each type (*dia*, *ELMO*, *Kinesin5* and *Map60* mutants, wild type). The nuclear displacement curves (in the first phase) are similar to the oscillation of a not-actively driven and non-damped harmonic oscillator. Therefore, the individual nuclear displacement curves were fitted to a sine curve of the form *y*(*t*) = *A* sin(*ωt* + *φ*), where *A* is the amplitude, *ω* the angular frequency and *φ* the phase shift, using a self-written script in Python. The biological rational behind this approach is that the nuclei behave like they were linked to an elastic spring, which could be e.g. linkages to the cytoskeleton. At t=0 the spring is stretched and the nuclei start to move until the spring is compressed and the nuclei move back.

The fit region was determined as follows. For all curves, the lower bound on the fitting range was set equal to the point in time where the nuclear displacement first exceeds 5 μm, as some curves show a small, reversible displacement in the beginning. The upper bound was chosen independently for each of the data sets since the elastic part of the curve depends on the stiffness and dampening of the spring and hence differs across data sets. For the *dia* and *Map60* mutants as well as the wild type, the upper bound was set equal to 100 s after the turning point, while it was set to 180 s after for the *ELMO* mutant and 240 s for the Kinesin5 mutants. The results of the fit parameter *ω* were averaged for each embryo to give a set of angular frequencies *ω*_*i*_ for each type, where *i* runs over the number of embryos. The spring constant was derived from the average *ω*_*i*_ *via* the relation *k* = *m* (*ω*_*i*_)^2^, in which *m* denotes the mass of the nucleus. It was assumed that the nuclei are spherical with a diameter of 4.9 μm and a density equal to that of water at room temperature. Error bars, which correspond to one standard deviation, were calculated in the frequency domain and then converted to the force domain by the analogue of the relation above.

## Acknowledgements

We are grateful to D. Görlich, T. Lecuit, B. Loppin and the members of the Grosshans laboratory for stains, materials and discussions. We acknowledge service support from the Bloomington Drosophila Stock Center (supported by NIH P40OD018537). X.Z. was supported by Center for Advancing Electronics Dresden (cfaed) and by the Deutsche Forschungsgemeinschaft (DFG, German Research Foundation) under Germany’s Excellence Strategy – EXC-2068 – 390729961– Cluster of Excellence Physics of Life of TU Dresden. This work was in part supported by the Göttingen Centre for Molecular Biology (funds for equipment repair) and the Deutsche Forschungsgemeinschaft (DFG, SFB-937/A10, SFB-937/A19 and equipment grant INST1525/16-1 FUGG).

## Author contributions

ZL and DK conducted the experiments. ZL, JR, HP, SK analyzed the data. SM, XZ, KA conducted the simulations. JG and ZL conceived and JG, TA, SG, KA supervised the study. JG, ZL and XZ wrote the manuscript.

---

**Figure S1.**
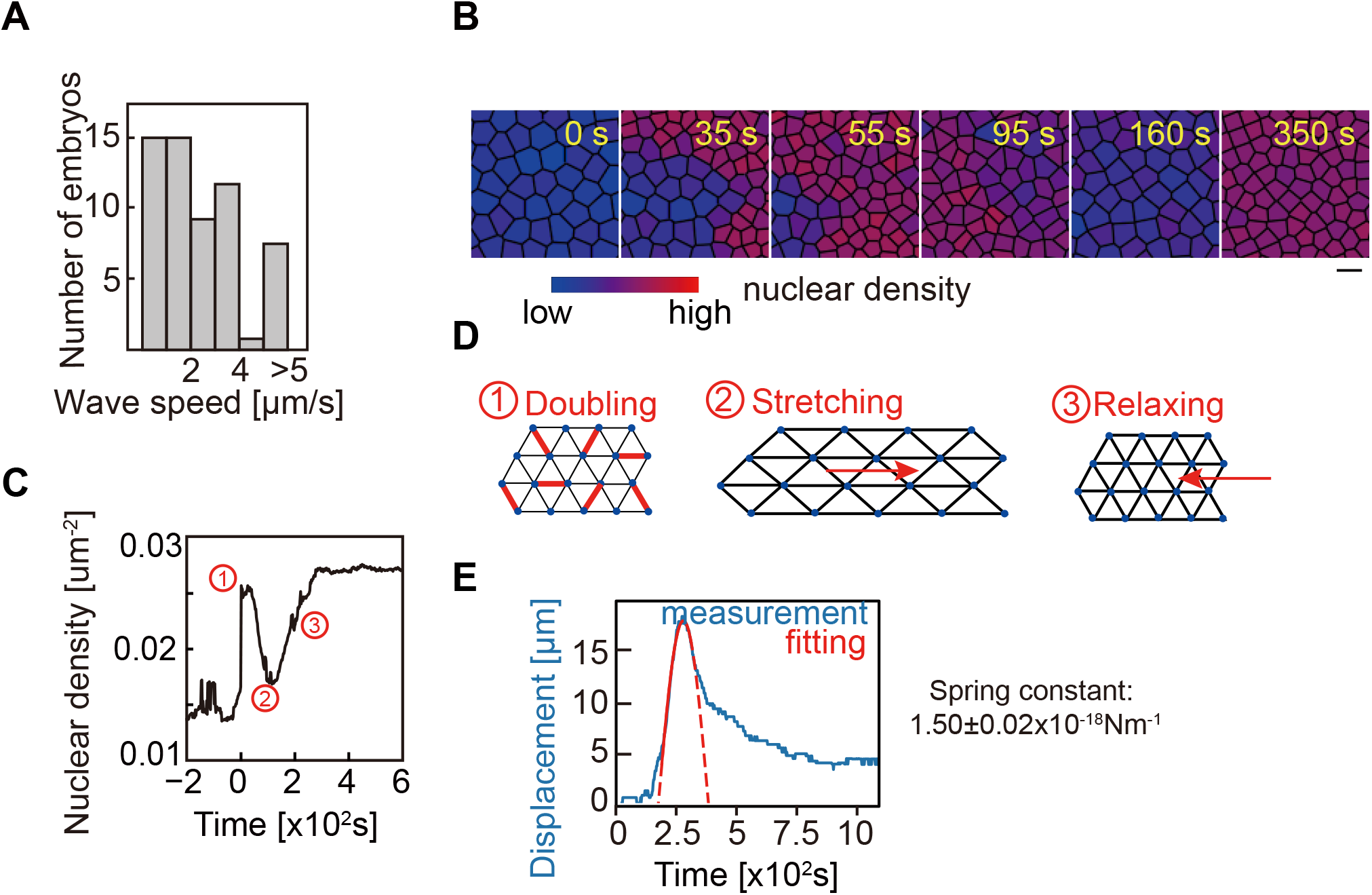
Characterization of nuclear dynamics in syncytial embryo. (A), Embryo to embryo variation of the speed of the wave front (n=50 embryos). (B), Image series with Voronoi maps with the mitotic wave front in images at 35 s and 55 s. Color code indicates nuclear density. (C), Time course of nuclear density with metaphase-anaphase transition at t=0. Numbers indicate the three stages of nuclear movement. (n=260 nuclei in one embryo). (D). Schematic drawing shows nuclei move like an elastic sheet. (E), A simple square function (red) was fitted to the forth and back movement around the maximal displacement. According to Hook’s law an apparent spring constant (1.5±0.02×10^−18^ Nm^−1^) was calculated (n=15 nuclei from 3 embryos). Data are mean±s.e.m. Scale bar: 10 μm.

**Figure S2.**
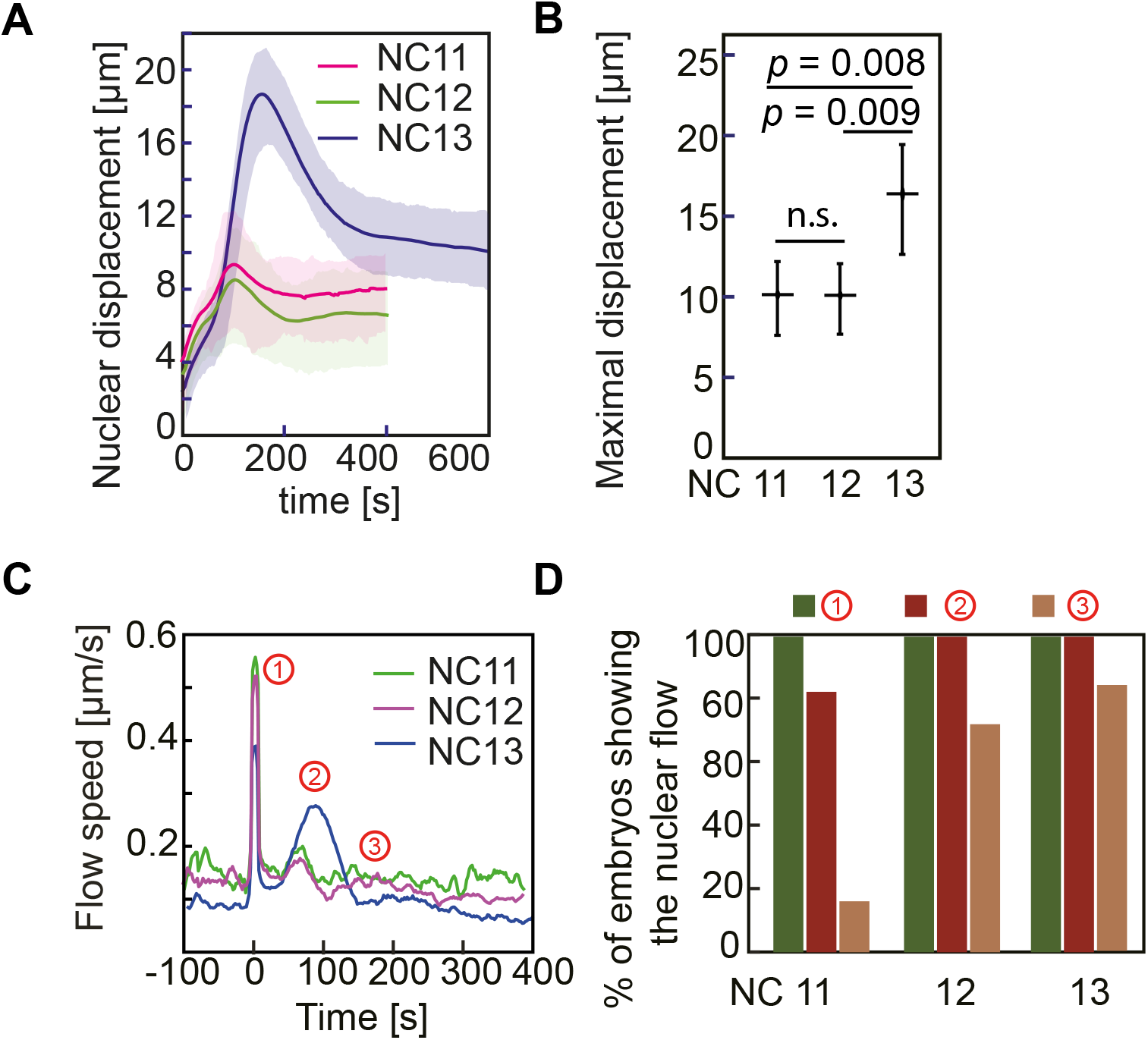
Nuclear displacement and flow speeds are less pronounced in earlier cycles. (A), The time course of nuclear displacement in NC11, 12 and 13 in one embryo (n=58, 107 and 206 nuclei in NC11, 12 and 13, respectively). (B), Maximal displacement distribution in NC11, 12 and 13 from (A). (C), Time course of nuclear flow speed from the same embryo as shown in a, with colour-coding for indicated nuclear cycles. (D), The percentage of embryos showing the nuclear movements ①, ② and ③ in different cycles (n=6, 18, 26 embryos in NC11, 12 and 13, respectively). Data are mean±s.e.m.

**Figure S3.**
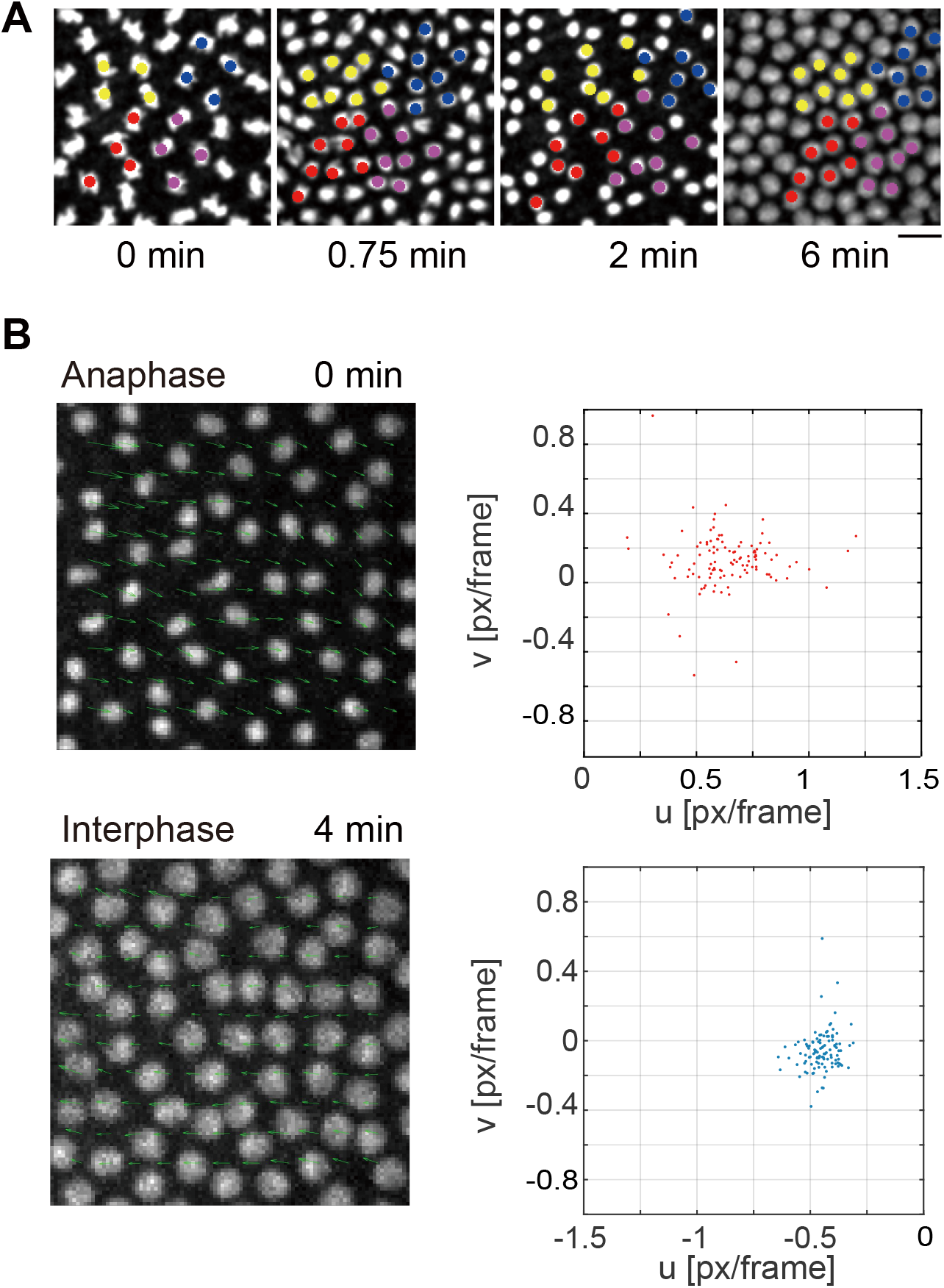
The nuclei move as a sheet. (A), Groups of cells have been marked in color at metaphase. Their daughter nuclei were labelled with the same color in the following images. (B) Particle Imaging Velocimetry analysis show that the nuclei move collectively like a sheet.

**Figure S4.**
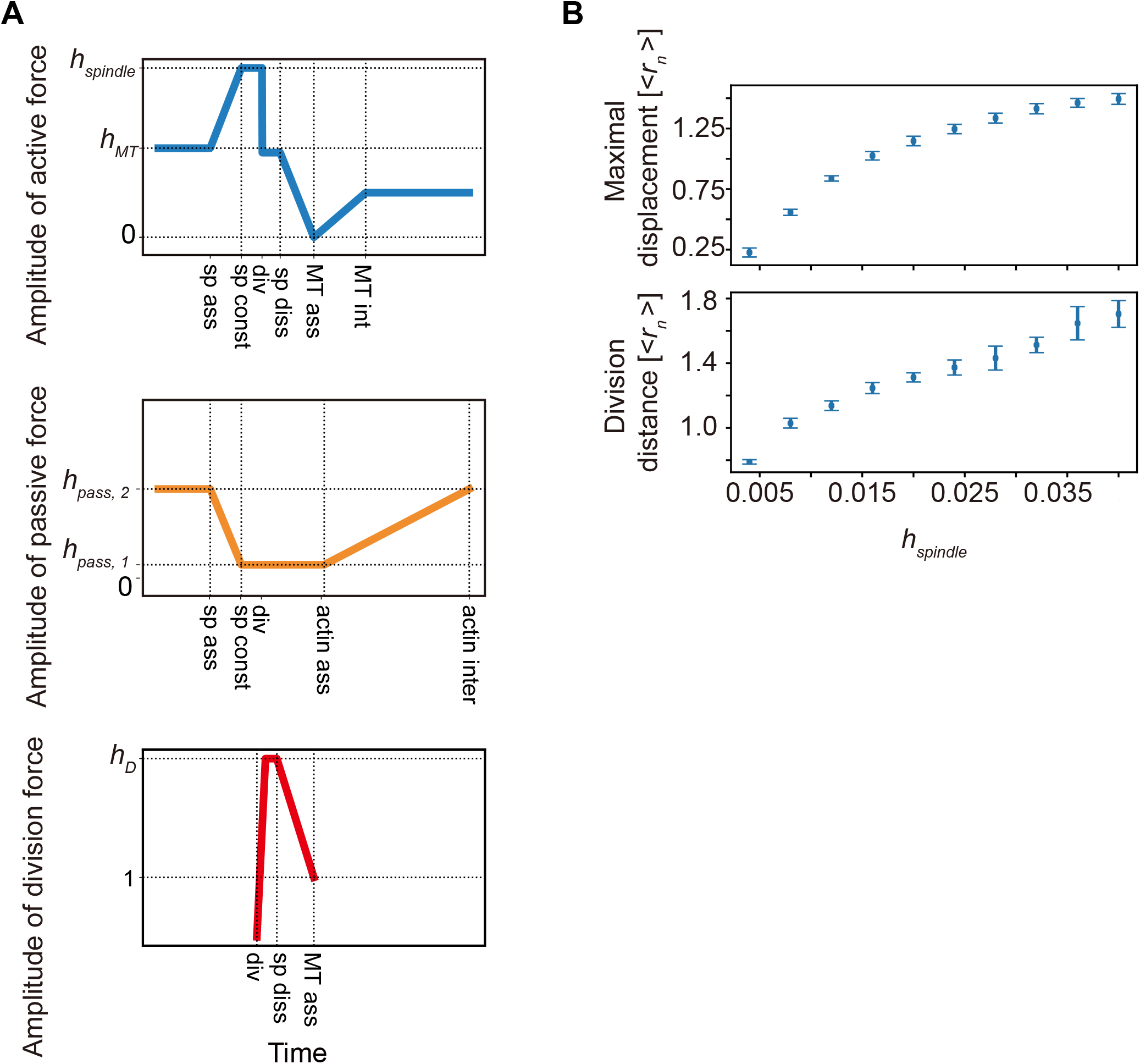
The parameters for Numerical simulation used in Fig. 3. (A), Time course of the division, active force and passive force used in the simulation. (B), Increasing in spindle strength (*h*_*spindle*_) leads to increasing of maximal displacement and division distance. Data are mean±s.e.m.

**Figure S5.**
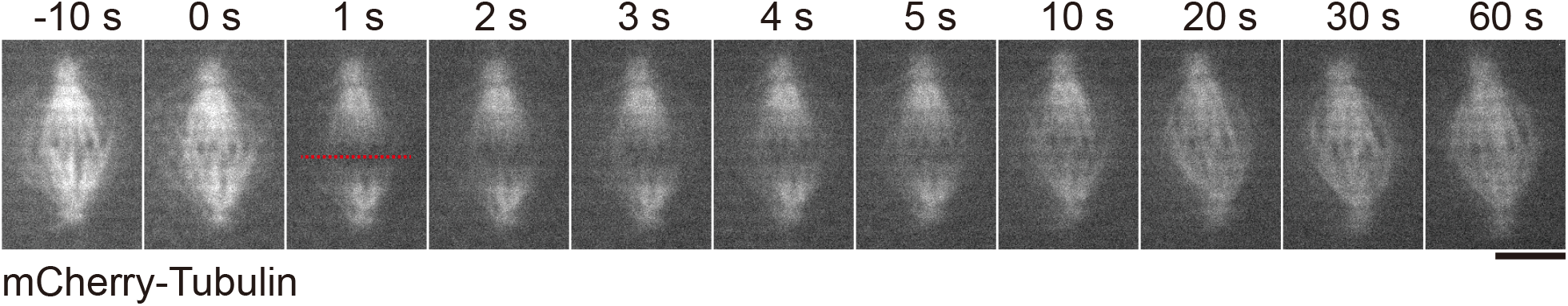
Laser cutting on spindle in metaphase. The embryo expressing mCherry-Tubulin was used for monitor the spindle structure. The spindle recovers in second-scale after laser ablation.

**Figure S6.**
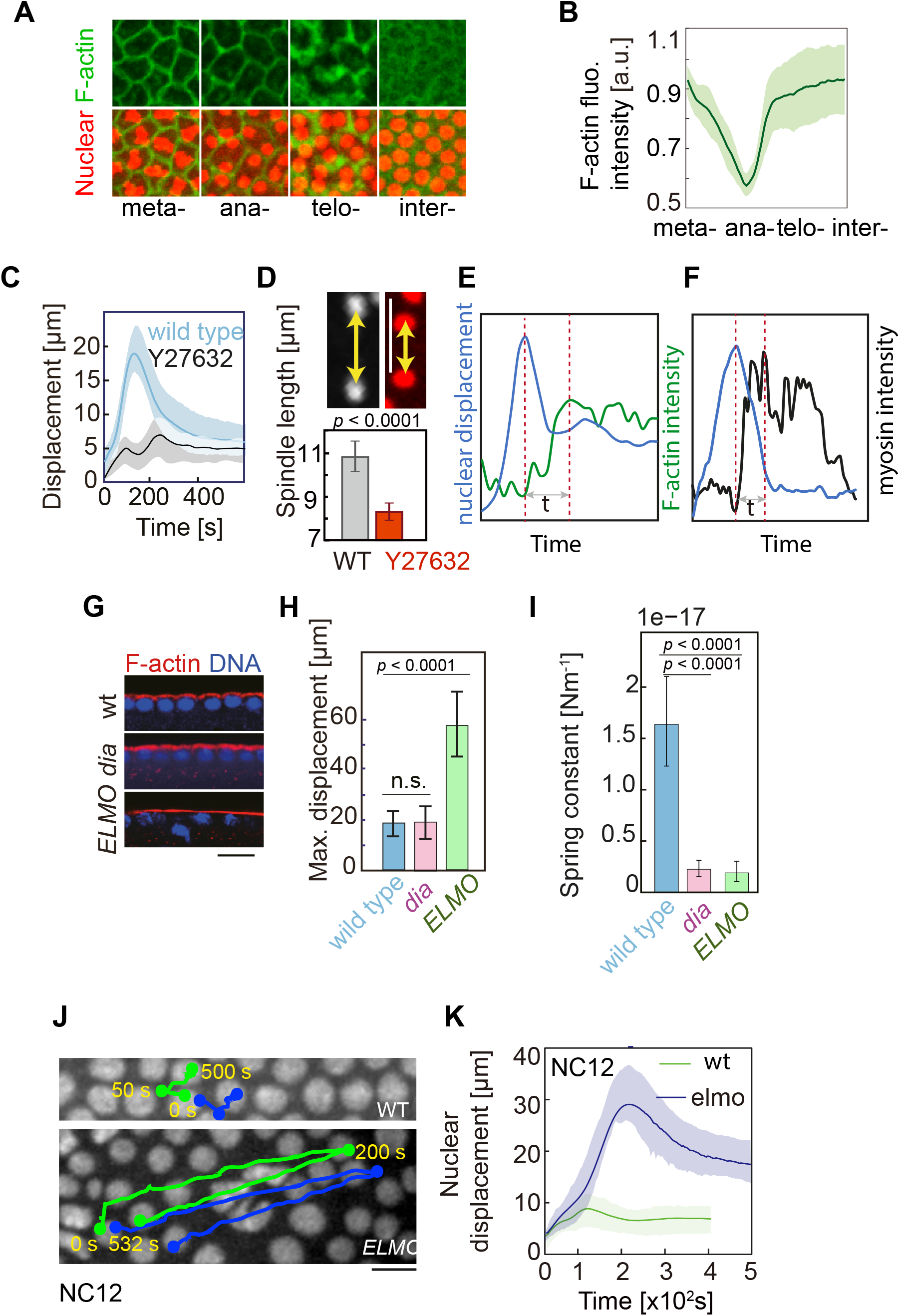
Actin cortex represses nuclear movement. (A), Live-images showing F-actin organization during nuclear cycle. (B), Quantification of F-actin. (n=10 regions in 3 independent recordings). (C, D), Injection of ROCK inhibitor Y-27632 leads to the reduction of nuclear motion as well as spindle elongation. (E, F), Time course of single nuclear displacement and the local myosin/F-actin intensity. (E), Fixed *dia* and *ELMO* mutants stained for F-actin (red) and DNA (blue). (F), Maximal displacement and final distance of nuclear dynamics in *dia* and *ELMO* mutants. (G), Apparent spring constant. (H), Images from movies of wild type and *ELMO* mutants in NC12. Nuclear trajectories are plotted into the first images. (I), The time course of nuclear displacement in NC12. (n=130 nuclei in *ELMO* and 290 nuclei in wild type. Data are mean±s.e.m. Scale bar: 10 μm.

**Figure S7.**
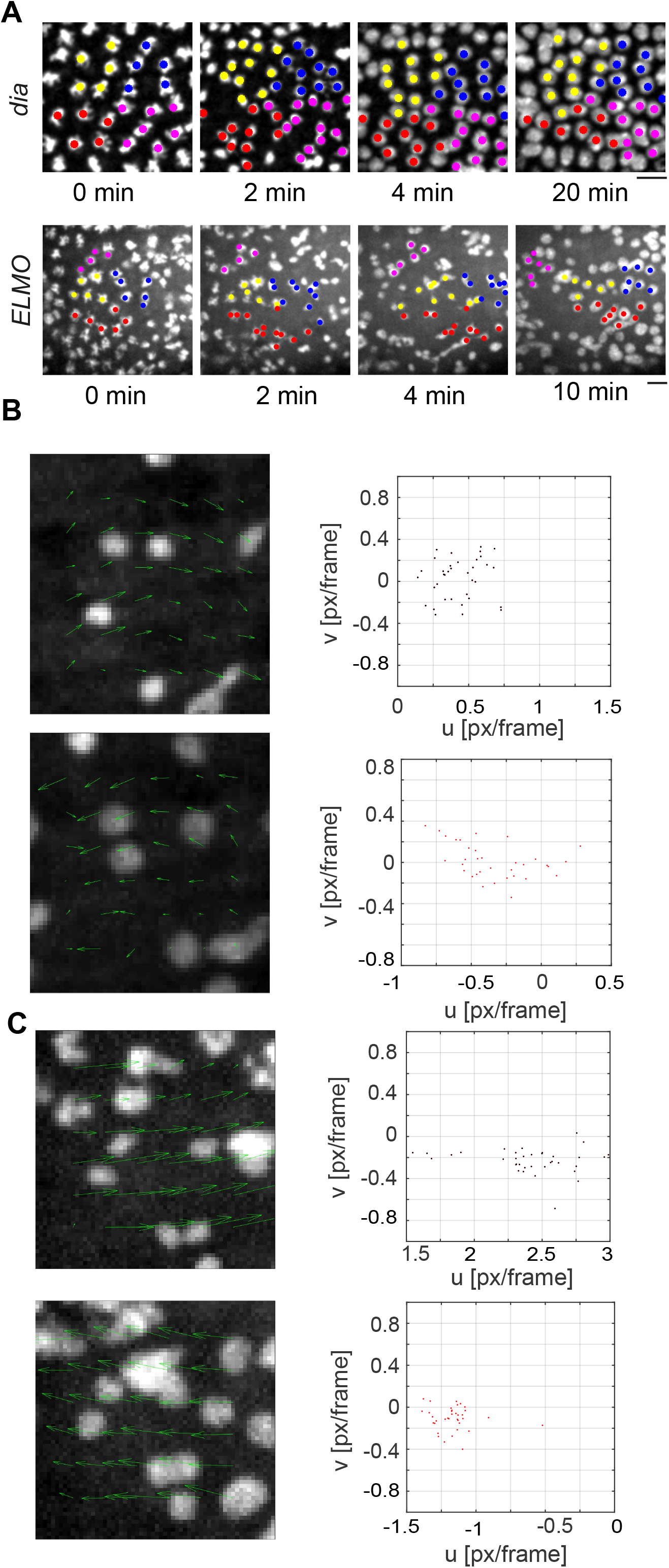
Nuclei move as a sheet in *dia* and *ELMO* mutant embryo. (A), Snapshots from movies at indicated time. Groups of cells have been marked in color at metaphase. Their daughter nuclei were labelled with the same color in the following images. (B). Particle Imaging Velocimetry analysis show that the nuclei move collectively.

**Figure S8.**
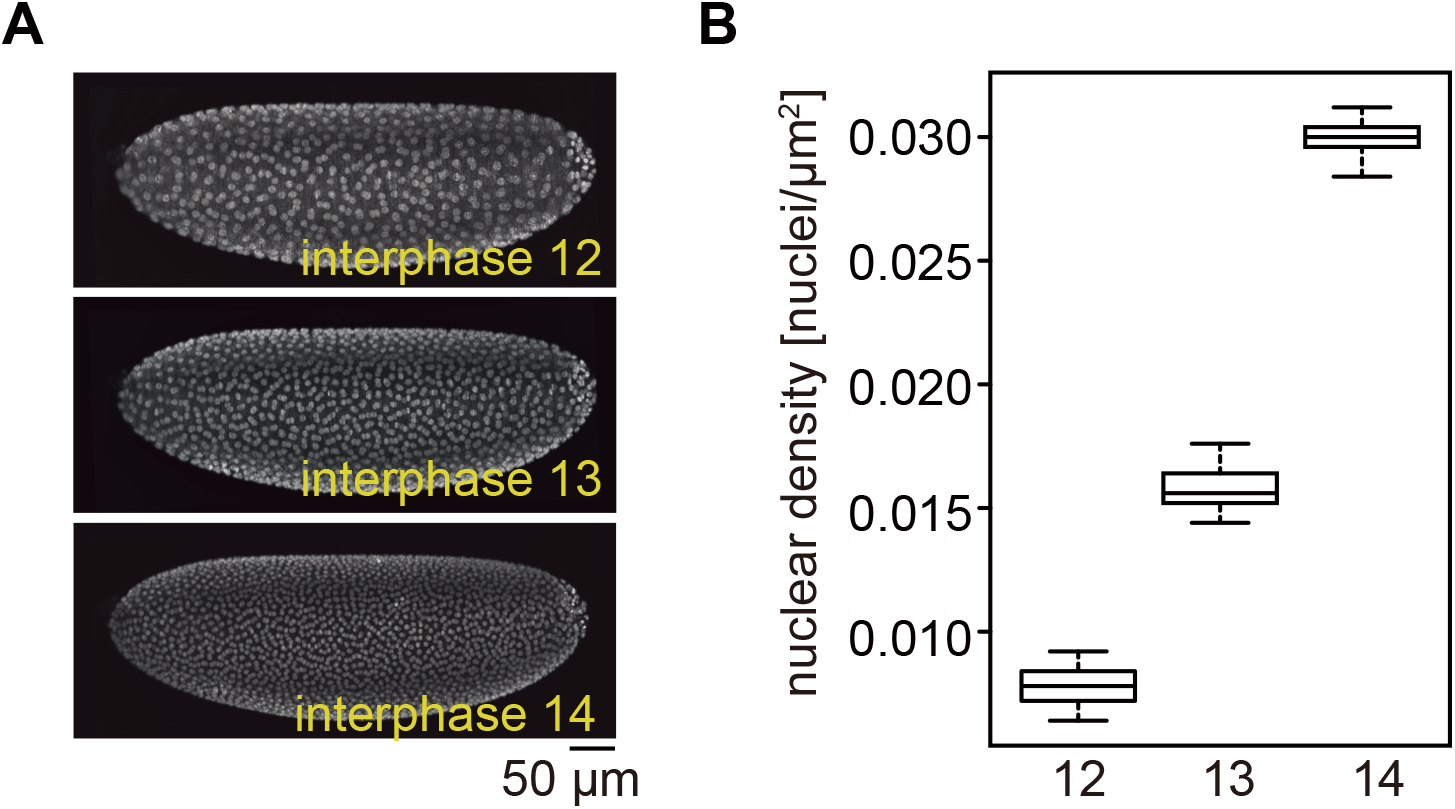
Nuclear densities in different cycles. (A), Images of Drosophila syncytial embryo expressing Histone2Av-GFP in different cycles. (B), Quantification of nuclear density in the indicated cycles. n=10 embryos. Data are mean±s.e.m.

**Figure S9.**
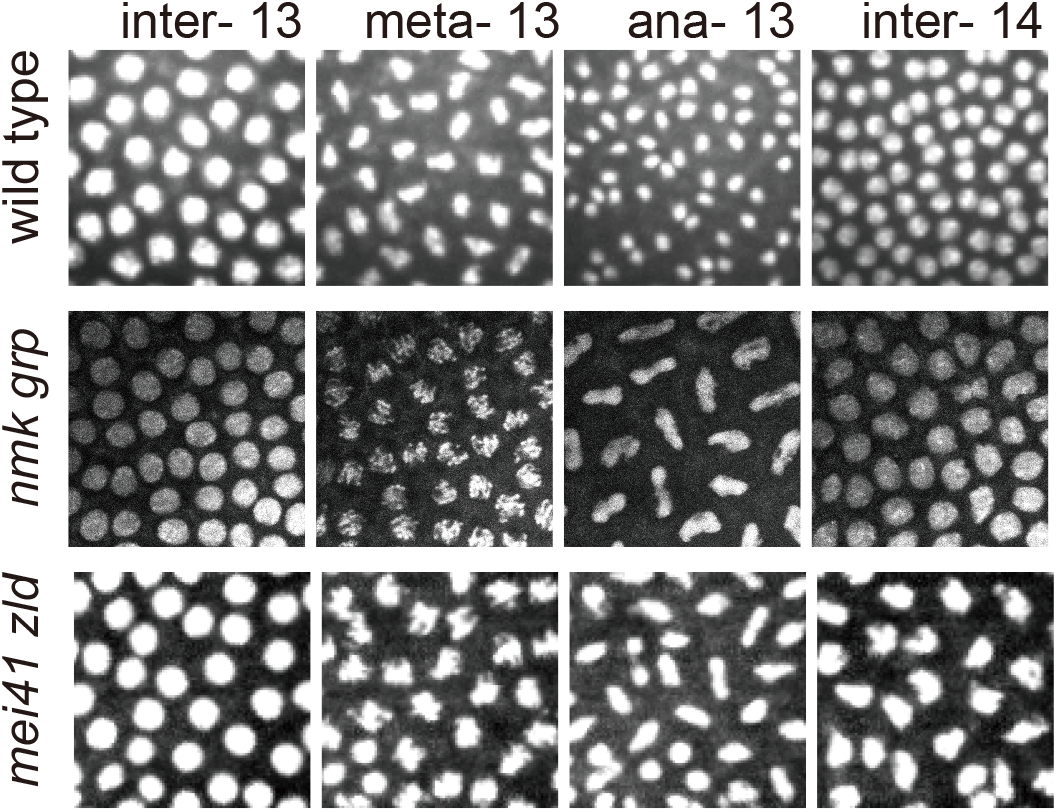
Daughter nuclei cannot separate in *nmk grp* and *mei41 zld* mutant embryos.

**Figure S10.**
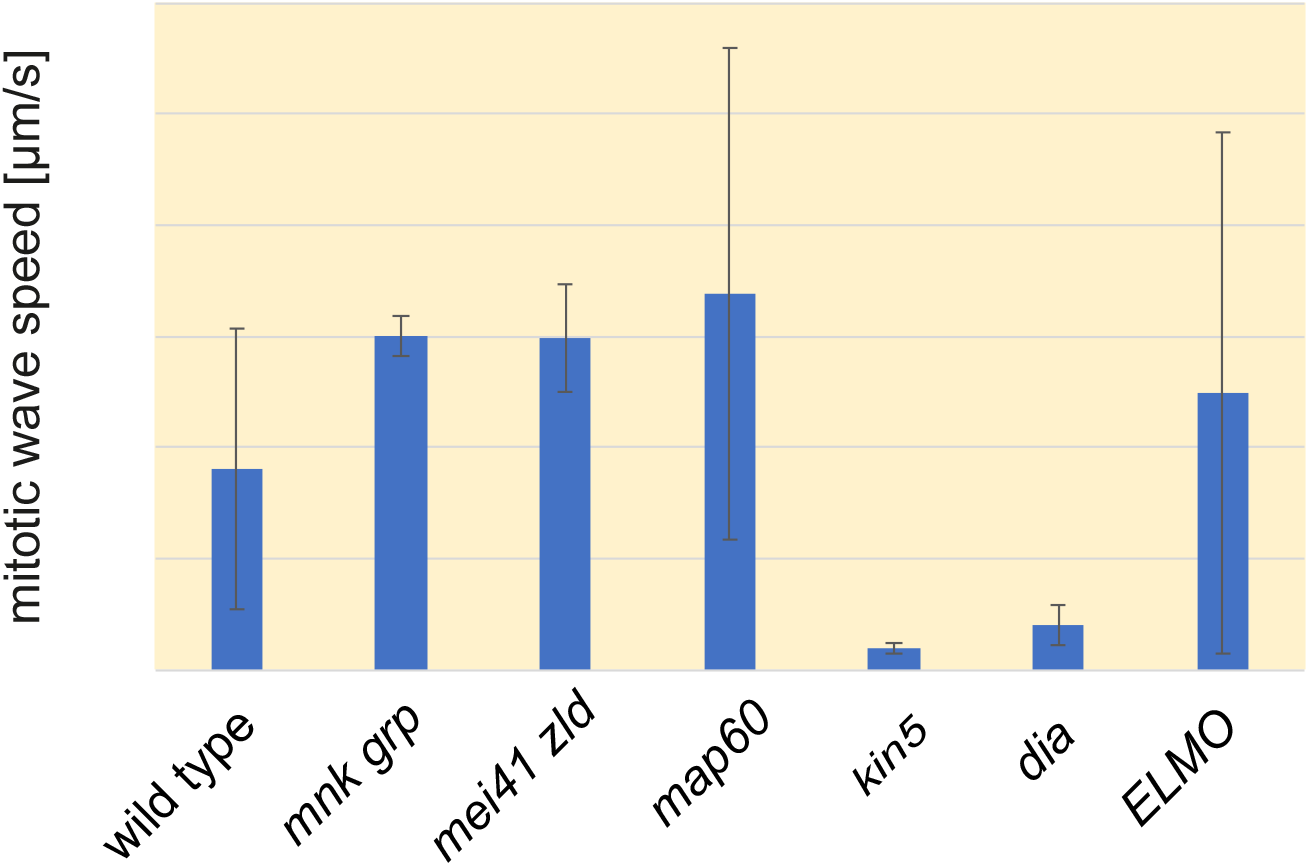
Mitotic wave speed in the indicated embryos.

**Figure S11.**
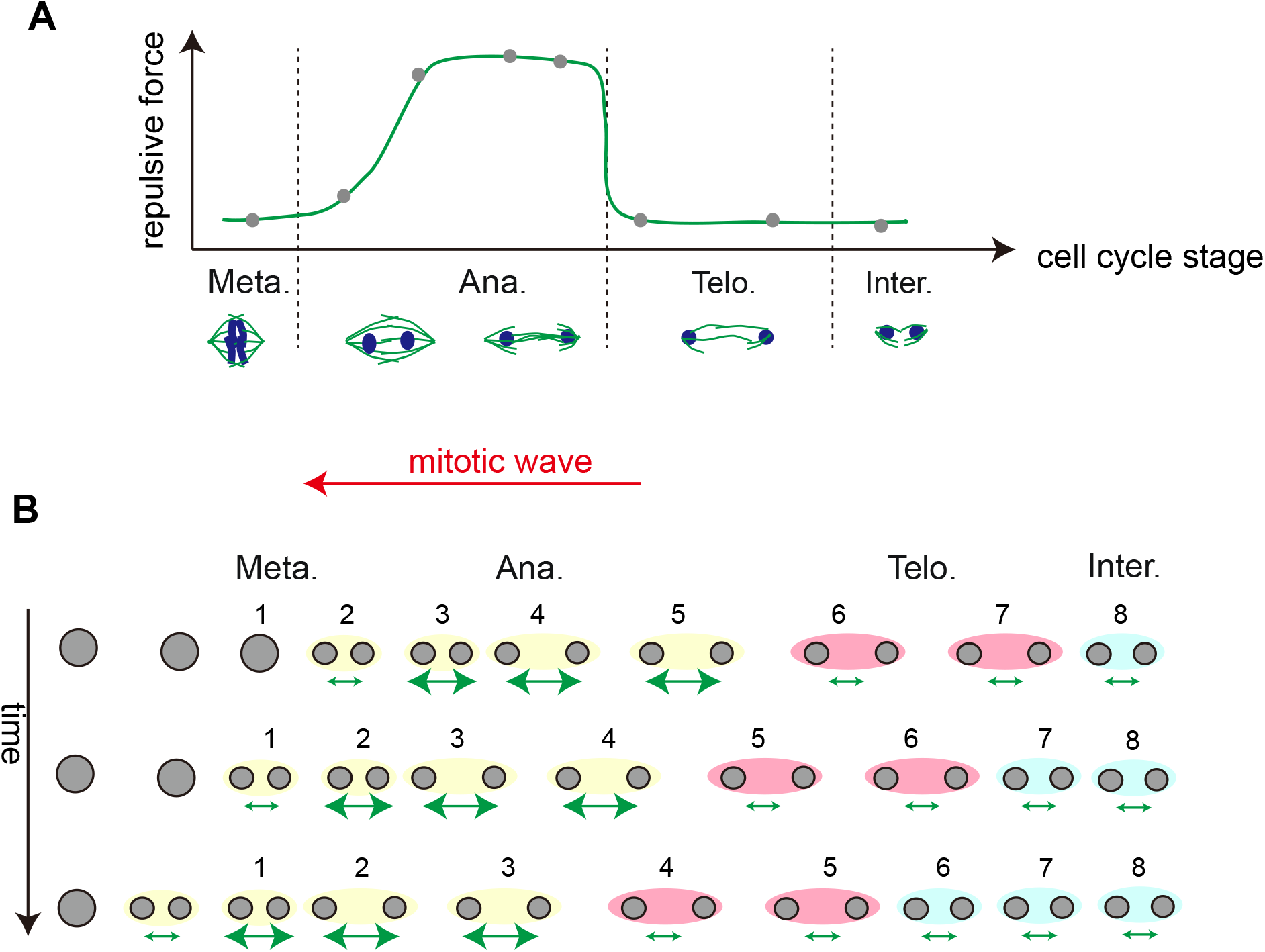
The asymmetric force field leads to the nuclear directional movement. (A), The repulsive force between daughter nuclei increases in anaphase pushing the daughter nuclei apart, followed by a drop in telophase due to spindle disassembly. (B), The summing up of all nuclei in an embryo at a given mitotic time results in an asymmetric force field, which likely determines the directionality of the nuclear flow in telophase.

## Movie list

Movie 1. Mitotic wave sweeps over the embryo.

Movie 2. Nuclei undergo stereotypical movement after metaphase-anaphase transition. Scale bar: 10 μm.

Movie 3. The time course of the nuclear Voronoi map over nuclear division. Scale bar: 10 μm.

Movie 4. Nuclei move as a laminar flow shown in PIV analysis. Scale bar: 10 μm.

Movie 5. Computational simulation.

Movie 6. The nuclear displacement in wild type, dia and ELMO mutant embryos. Scale bar: 20 μm.

